# One-pot Detection of COVID-19 with Real-time Reverse-transcription Loop-mediated Isothermal Amplification (RT-LAMP) Assay and Visual RT-LAMP Assay

**DOI:** 10.1101/2020.04.21.052530

**Authors:** Deguo Wang

## Abstract

**Background:** Rapid and reliable diagnostic assays were critical for prevention and control of the coronavirus pneumonia caused by COVID-19.

**Objective:** This study was to establish one-pot real-time reverse-transcription loop-mediated isothermal amplification (RT-LAMP) assay and one-pot visual RT-LAMP assay for the detection of COVID-19.

**Methods:** Six specific LAMP primers targeting the N gene of COVID-19 were designed, the RT-LAMP reaction system was optimized with plasmid pUC57 containing N gene sequence, the detection limit was determined with a serial dilution of the plasmid pUC57 containing N gene sequence, and the one-pot real-time RT-LAMP assay and one-pot visual RT-LAMP assay for the detection of COVID-19 were established.

**Results:** Our results showed that the one-pot RT-LAMP assays can detect COVID-19 with a limit of ≥ 6 copies per *μ*l^−1^ of pUC57 containing N gene sequence.

**Conclusion:** This study provides rapid, reliable and sensitive tools for facilitating preliminary and cost-effective prevention and control of COVID-19.

## Introduction

The outbreak of COVID-19, firstly reported from Chinese Wuhan in December 2019, has spread to many other countries in the last few weeks^1^, and has caused almost 80 000 infections and more than 2000 people died. The virus spreads so quickly that it has attracted the globe attention, and rapid diagnosis is one of the effective ways to prevent and control the diseases.

Chest CT and RT-PCR has been used for the clinical diagnosis of the coronavirus pneumonia^2, 3^, however, chest CT needs CT equipment and put the professional operator at risk of infection, and RT-PCR has high professional and technical requirements for operators.

Loop-mediated isothermal amplification (LAMP) can amplify nucleic acids under isothermal conditions^4^, and it had the advantages over real-time PCR assays in specificity sensitivity, cost effectiveness and rapidity^5, 6, 7^. The objective of this study was to develop the one-pot real-time RT-LAMP assay and the one-pot visual RT-LAMP assay for diagnosis of the pneumonia caused by COVID-19, which would be suitable for use at less developed areas.

## Materials and Methods

### Primer design for RT-LAMP assays

The RT-LAMP primers targeting the N gene of COVID-19 (GenBank accession No. MN997409.1) were designed using PrimerExplorer V5 (http://primerexplorer.jp/e/) and Oligo 7 (Molecular Biology Insights, Inc. Colorado Springs, CO, USA) software packages. The primer sequences are GCC AAA AGG CTT CTA CGC A (F3), TTT GGC CTT GTT GTT GTT GG (B3), TCC CCT ACT GCT GCC TGG AGT TTT CGG CAG TCA AGC CTC TTC (FIP), TCC TGC TAG AAT GGC TGG CAA TTT TTT TTG CTC TCA AGC TGG TTC A (BIP), CGA CTA CGT GAT GAG GAA CGA (LF) and GCG GTG ATG CTG CTC T (LB), Table 1, and the length of the targeted sequence was 233 bp.

### Preparation of nucleic acid samples

The N gene (GenBank accession No. MN997409.1) of COVID was chemically synthesized and cloned into pUC57 plasmid (herein referred to as pUC57-N DNA) by General Biosystems (Anhui) Co., Ltd, the pUC57-N DNA was used as the template for optimization of the RT-LAMP system, as well as for determination of sensitivity.

### Optimization of the real-time RT-LAMP reaction remperature

The real-time RT-LAMP assay with above designed RT-LAMP primers was performed in a 50-μL reaction mixture containing 0.8 mM each of forward inner primer (FIP) and backward inner primer (BIP), 0.2 mM each of forward outer primer (F3) and backward outer primer (B3), 0.4 mM of forward loop primer (LF) and backward loop primer (LB), 1.2 mM dNTPs, 1× Bst DNA Polymerase Buffer (Zhengzhou Shenxiang Industrial Co., Ltd, China), 1 × EvaGreen, 1 × Rox, 1 pg pUC57-N DNA, and 25 U Bst DNA/RNA Polymerase 3.0 (New England Biolabs, Inc., MA, USA) ^8, 9^. The reaction mixtures were heated at 55°C, 57°C, 59°C and 61°C for 50 min (30 s per cycle), individually. The amplification plot and melt curve were obtained using a StepOne™ System (Applied Biosystems, Foster City, CA, USA).

### Effect of inhibitors in blood samples on amplification efficiency of RT-LAMP assays

Because the RNA extraction step was omitted, the effect of inhibitors in blood samples on the amplification efficiency of the RT-LAMP assays was to be determined. 10 µL, 7.5µL, 5 µL and 2.5 µL pork blood samples (purchased from Xuchang Market, China) free of COVID-19 were dissolved in 10 µL 2019-nCoV-Fast-Sample Nucleic Acid Releasing Agent, respectively, which were added to above 50 µL real-time RT-LAMP reaction system and heated at the optimized temperature for 50 min (30 s per cycle) in StepOne™ System (Applied Biosystems, Foster City, CA, USA).

### Sensitivity Determination of the Real-Time RT-LAMP Assay and the Visual RT-LAMP Assay

The analytic sensitivities of the newly developed real-time RT-LAMP assay and visual RT-LAMP assay were determined with pUC57-N DNA ranging from 2-0.02 fg, and the reaction mixtures were heated at the optimal temperature for 60 min in a StepOneTM System (Applied Biosystems, Foster City, CA, USA) or in a water bath, when water bath was used, the reaction tube was sunk into water. The 50-μL reaction mixture contained 0.8 mM each of forward inner primer (FIP) and backward inner primer (BIP), 0.2 mM each of forward outer primer (F3) and backward outer primer (B3), 0.4 mM of forward loop primer (LF) and backward loop primer (LB), 1.2 mM dNTPs, 1× Bst DNA Polymerase Buffer (Zhengzhou Shenxiang Industrial Co., Ltd, China), 0.1 mM MnSO_4_, 0.1 mM 4-(2-pyridylazo) resorcinol, 5 µL pork blood samples free of COVID-19, 10 µL 2019-nCoV-Fast-Sample Nucleic Acid Releasing Agent (Hunan Shengxiang Biology Technology Co., Ltd, China), 25 U Bst DNA/RNA Polymerase 3.0 (New England Biolabs, Inc., MA, USA), and pUC57-N DNA of 2-0.02 fg^9^.

### Stability and Repeatability Tests of the Real-Time LAMP Assay

The stability and the repeatability of the real-time LAMP system were tested with one-month interval. The test was carried out at the optimized temperature determined as described above with four positive controls (1 pg pUC57-N DNA) and four negative controls (ddH_2_O) in a StepOne™ System (Applied Biosystems, Foster City, CA, USA).

## Results and Analysis

### Optimization of the real-time RT-LAMP reaction temperature

For the RT-LAMP reaction, as shown in Figure 1, all positive controls (with pUC57-N DNA as template) had amplification, and the results of all negative controls (DNA template substituted by ddH_2_O) were negative at 55°C, 57 °C, 59 °C and 91 °C, however, there was no significant difference in the amplification efficiency, and 59°C was temporarily was selected for the subsequent experiments.

**Figure 1.**
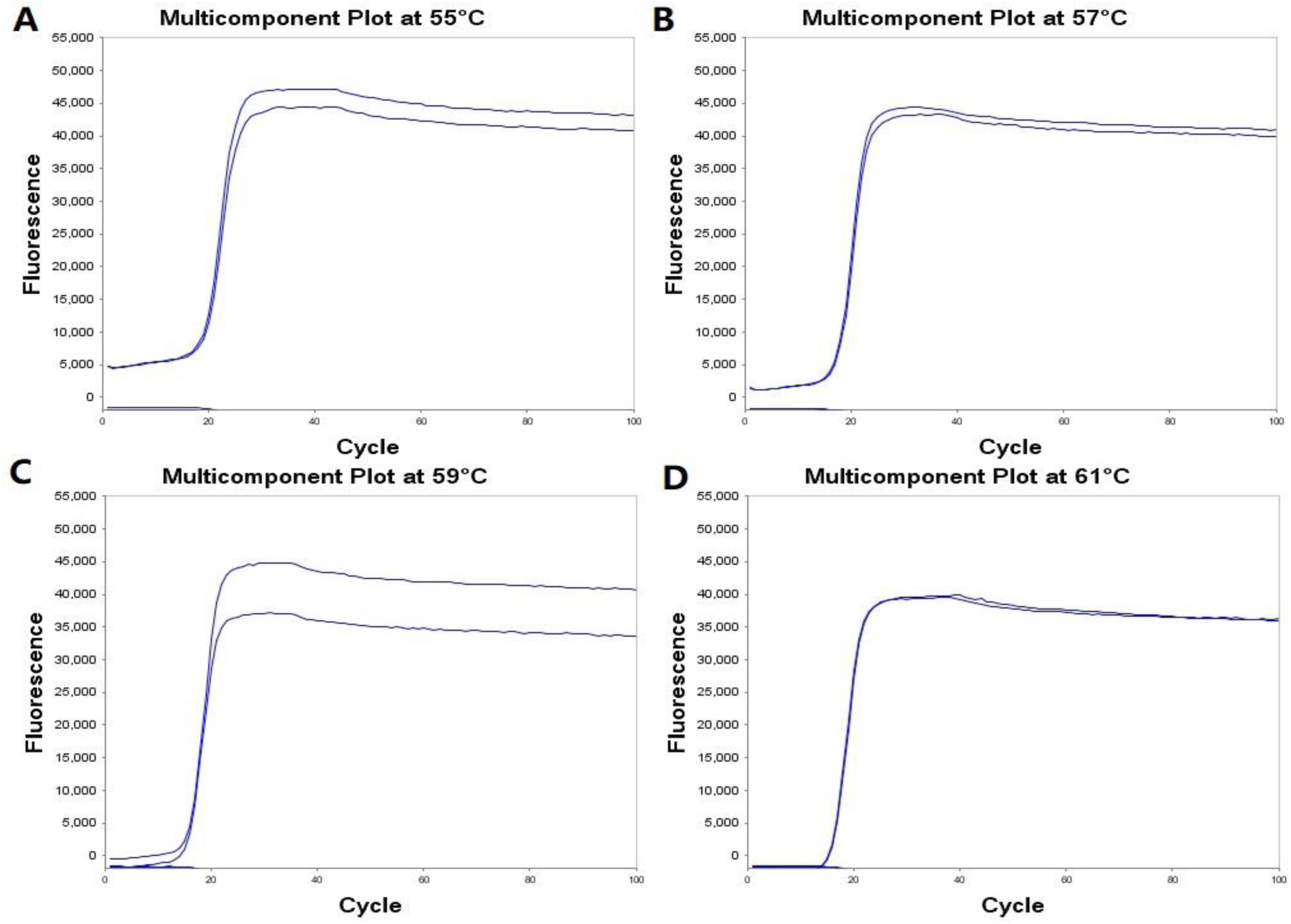
Amplification Plots of the Real-Time RT-LAMP Reactions. A: heated at 55°C for 50 min; A: heated at 57°C for 50 min; A: heated at 59°C for 50 min; A: heated at 61°C for 50 min.

### Effect of inhibitors in blood samples on the RT-LAMP amplification assays

The effect of inhibitors in blood samples on the amplification efficiency had been determined, as Figure 2 indicated, when the real-time RT-LAMP reaction system was added with 10 µL blood samples dissolved in 10 µL 2019 -nCoV-Fast-Sample Nucleic Acid Releasing Agent (Hunan Shengxiang Biology Technology Co., Ltd, China), there was no amplification; when 7.5 µL blood samples and 10 µL 2019-nCoV-Fast-Sample Nucleic Acid Releasing Agent added, there was slight amplification; when 5µL or 2.5 µL blood samples dissolved in 10 µL 2019-nCoV-Fast-Sample Nucleic Acid Releasing Agent added, there was acceptable effect on the real-time RT-LAMP reaction. Therefore, the blood sample volume of 5µL was selected for 50 µL one-pot real-time or visual RT-LAMP reaction.

**Figure 2.**
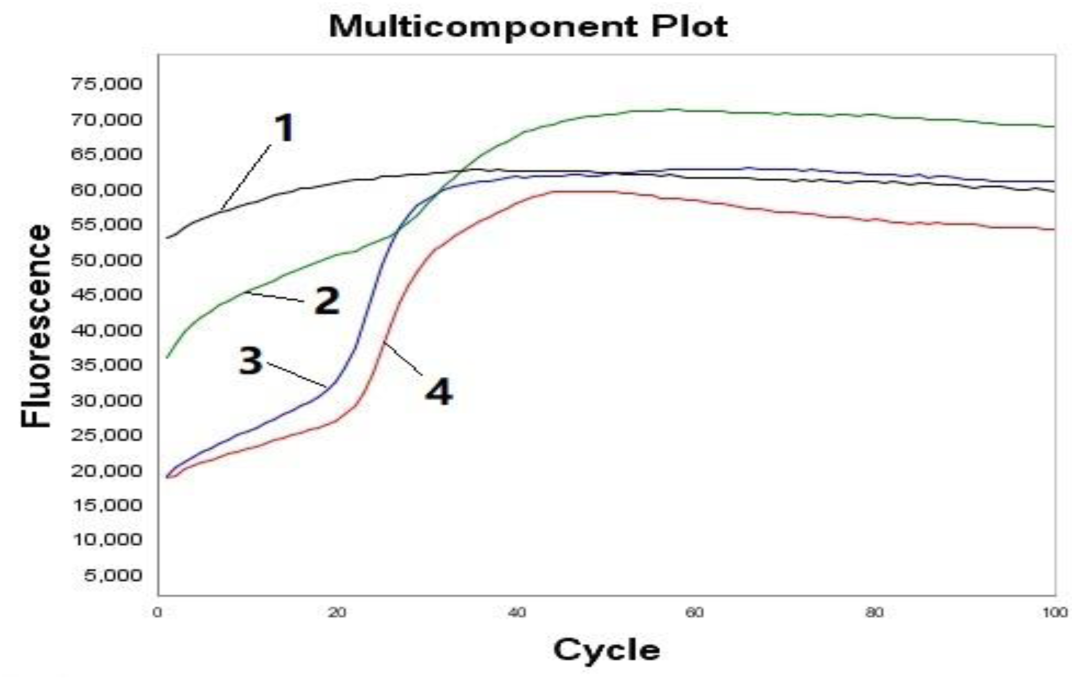
Effect of Inhibitors in Blood Samples on Amplification of Real-time RT-LAMP Assay. 1. 10 µL blood samples dissolved in 10 µL 2019-nCoV-Fast-Sample Nucleic Acid Releasing Agent was added to 50 µL the reaction system; 2. 7.5 µL blood samples dissolved in 10 µL 2019-nCoV-Fast-Sample Nucleic Acid Releasing Agent was added to 50 µL the reaction system; 3. 5 µL blood samples dissolved in 10 µL 2019-nCoV-Fast-Sample Nucleic Acid Releasing Agent was added to 50 µL the reaction system; 4. 2.5 µL blood samples dissolved in 10 µL 2019-nCoV-Fast-Sample Nucleic Acid Releasing Agent was added to 50 µL the reaction system.

### Sensitivity of the one-pot real-time or visual RT-LAMP assay

The detection limits of the one-pot real-time RT-LAMP assay and the one-pot visual RT-LAMP assay were determined using pUC57-N DNA ranging from 2-0.02 fg at 59 °C for 60 min in a StepOneTM System (Applied Biosystems, Foster City, CA, USA) or in a water bath. The detection limits of both the one-pot real-time RT-LAMP assay and the one-pot visual RT-LAMP assay were found to be ≥ 0.2 fg pUC57-N DNA (Figure 3), which was equivalent to ≥ 6 copies.

**Figure 3.**
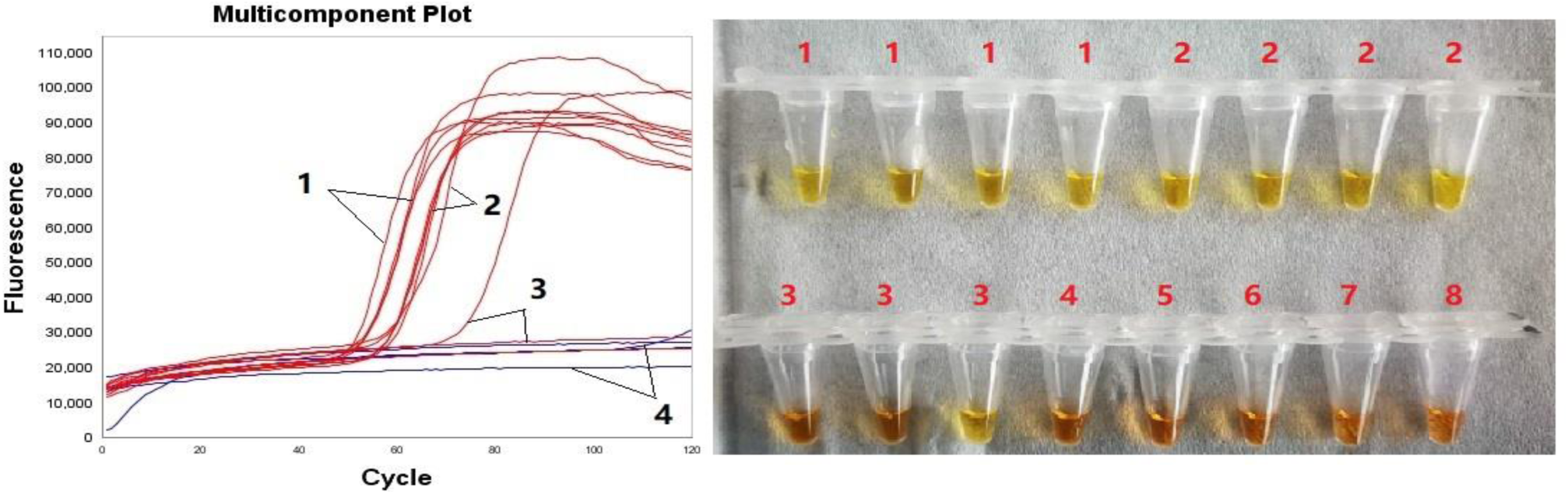
Sensitivity of the Developed One-pot Real-Time RT-LAMP Assay and One-pot Visual RT-LAMP Assay. 1. 2 fg pUC57-N DNA; 2. 0.2 fg pUC57-N DNA; 4. 0.02 fg pUC57-N DNA; 4. H_2_O (Negative Controls). Note: positive reaction: color changed from orange to light yellow; negative reaction: orange.

### Stability and Repeatability of the Real-Time LAMP Assay

The stability and the repeatability of the real-time LAMP system were tested with one-month interval. As shown in Figure 4, the reactions of four positive controls were positive with the same amplification plot and melt curve, while the reactions of all negative controls were negative with the same melt curve. This demonstrated that the newly established real-time LAMP assay was robust and repeatable.

**Figure 4.**
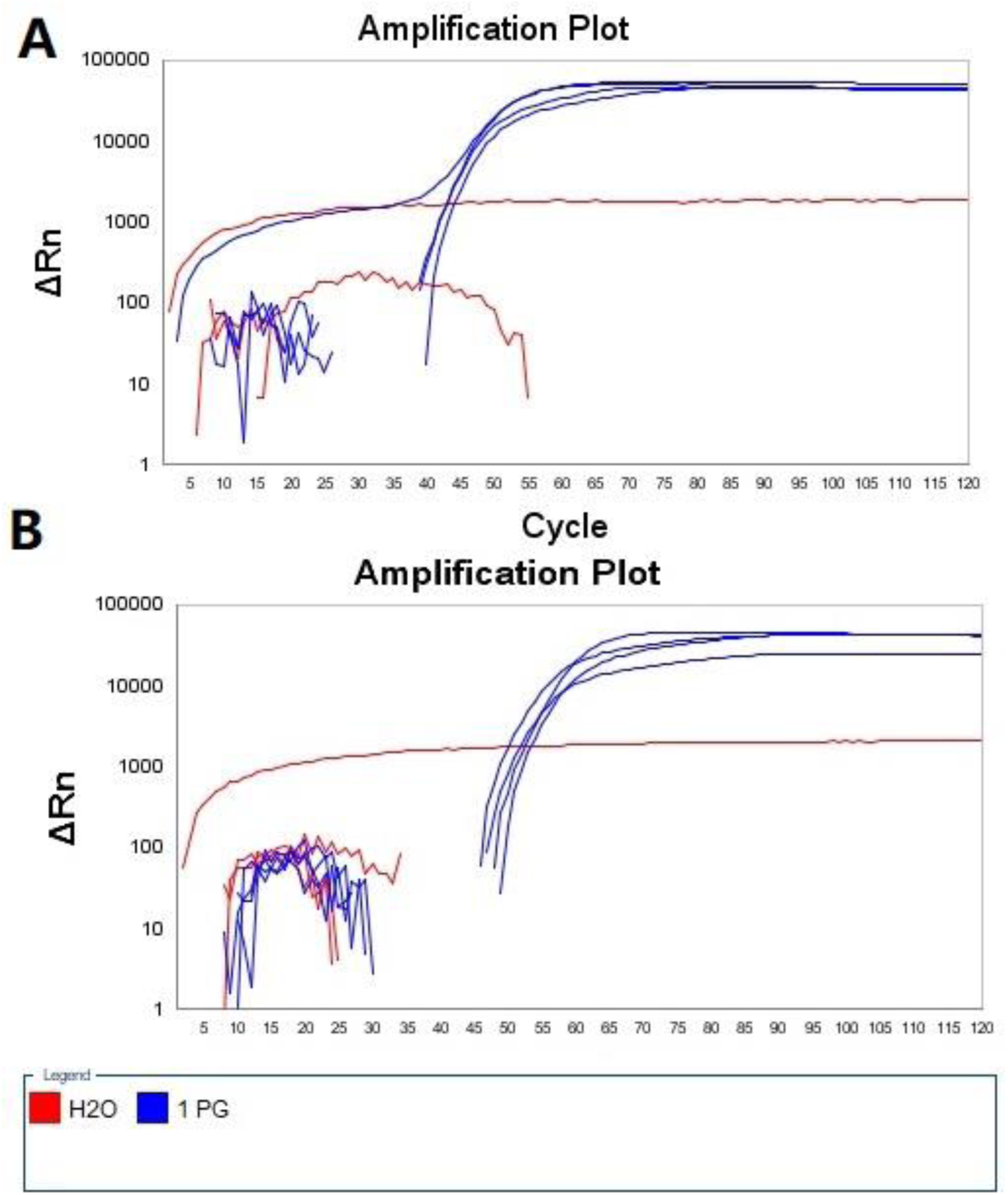
Repeatability of Newly Established Real-Time LAMP Assay. Note: eight positive controls with 1 pg pUC57-N DNA as the template; eight negative controls with ddH_2_O as the template; A: tested on Mar 19, 2020; B: Tested on Apr 20, 2020.

## Discussion

The one-pot real-time RT-LAMP assay and one-pot visual RT-LAMP assay for detection of COVID-19 were established in the study, and the detection limit was ≥ 6 copies. Although the specificity of the established RT-LAMP assays had been not determined, they were still considered to be highly specific for following reason. Upon alignment in DNA Data Bank of Japan, the target sequence of established RT-LAMP assays had 100% identities (233/233) with that of 35 COVID-19 strains, and had 88% identities (207/233) with that of 52 congeneric SARS Coronavirus strains, among which there were 12 bp on the RT-LAMP primers, it was reported that 3 bp primer-template mismatches extend the detection time from 21 min to 47 min^10^, therefore, the established RT-LAMP assays were theoretically highly specific.

The Bst DNA/RNA Polymerase 3.0 (New England Biolabs, Inc., MA, USA) had been used in the study. The enzyme has faborable perforance of both amplification and reverse transcription activity, so it can use either DNA or RNA as template ^11,12^, no reverse transcriptase is needed, and the estalished assays can be directly used for detection of the RNA virus.

The nucleic acid extraction had been omitted in the established one-pot real-time or visual RT-LAMP assays, which had greatly reduced the infection risk of the operators. Furthermore, it was recommended the visual RT-LAMP assay over the real-time RT-LAMP assay, one of four Negative Controls in above sensitivity determination of the real-time RT-LAMP assay had amplification, while all four Negative Controls of the visual RT-LAMP assay had no amplification, because the aerosol that may leak was washed by water in water bath, the false positive rate caused by aerosol of the real-time RT-LAMP assay was higher than that of the visual RT-LAMP assay which sunk the reaction tube in water.

## Conclusion

This study had established the one-pot real-time RT-LAMP assay and one-pot visual RT-LAMP assay for the detection of COVID-19, and provided the rapid, reliable and sensitive tools for facilitating preliminary and cost-effective prevention and control of COVID-19.

